## supplementary data for "Proteomics identifies substrates and a novel component in hSnd2-dependent ER protein targeting"

### Proteomics identifies substrates and a putative novel component in hSnd2-dependent protein targeting to the human ER

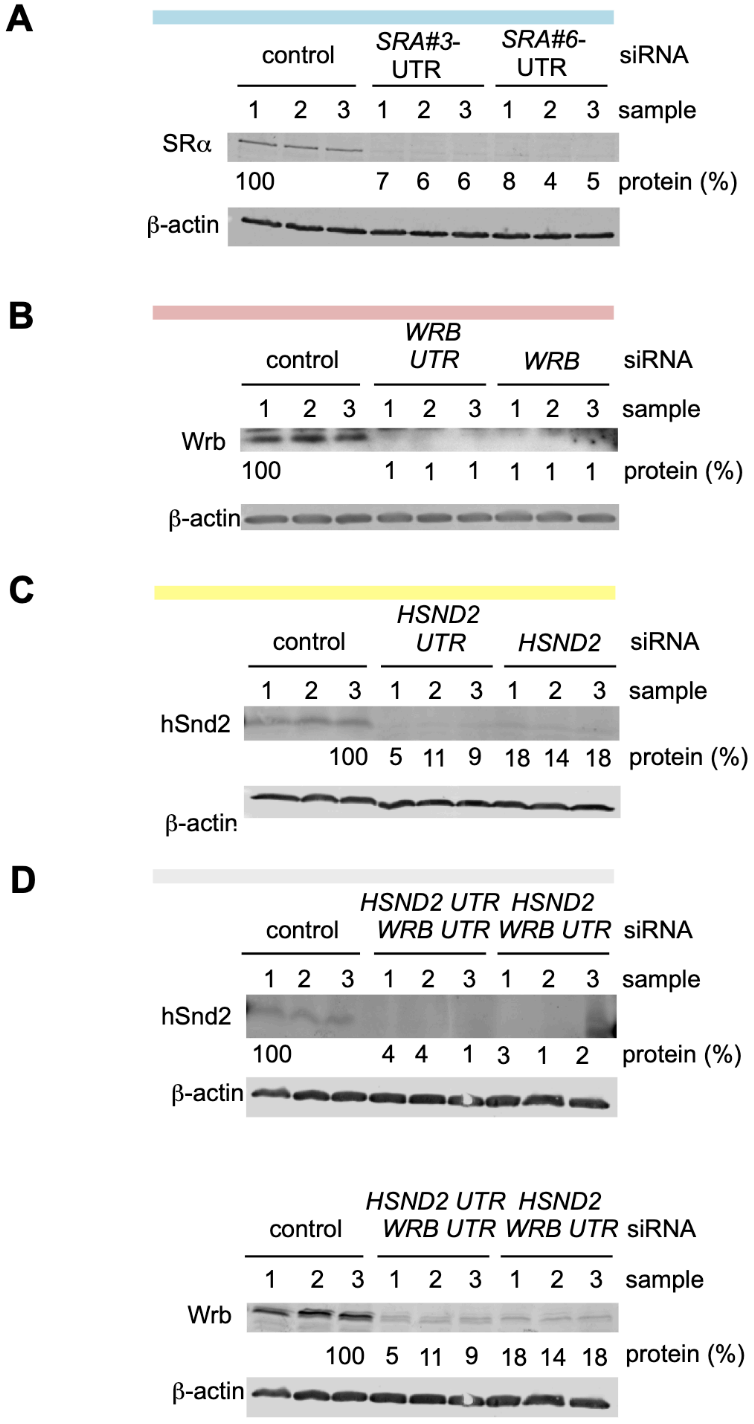
A
control
SRA#3-UTR
SRA#6-UTR
siRNA
1 2 3 1 2 3 1 2 3 sample
SRα
100 7 6 6 8 4 5 protein (%)
β-actin
B
control
WRB UTR
WRB
siRNA
1 2 3 1 2 3 1 2 3 sample
Wrb
100 1 1 1 1 1 1 protein (%)
β-actin
C
control
HSND2 UTR
HSND2
siRNA
1 2 3 1 2 3 1 2 3 sample
hSnd2
100 5 11 9 18 14 18 protein (%)
β-actin
D
control
HSND2 UTR WRB UTR
HSND2 WRB UTR
siRNA
1 2 3 1 2 3 1 2 3 sample
hSnd2
100 4 4 1 3 1 2 protein (%)
β-actin
control
HSND2 UTR WRB UTR
HSND2 WRB UTR
siRNA
1 2 3 1 2 3 1 2 3 sample
Wrb
100 5 11 9 18 14 18 protein (%)
β-actin

**Figure S1. Western blots confirming depletion of various components for protein targeting to the ER by quantitative MS in HeLa cells. Related to Figures 1 and 3.** (A-D) The experimental strategy in the MS experiments involved siRNA-mediated gene silencing using two different siRNAs for each target and one non-targeting (control) siRNA, respectively with three replicates for each siRNA, label-free quantitative proteomic analysis and differential protein abundance analysis to identify negatively affected proteins (i.e. clients) as well as positively affected proteins (i.e. compensatory mechanisms). Knock-down efficiencies were evaluated by Western blot. Only the respective areas of interest are shown. Results are presented as % of residual protein levels (normalized to ß-actin) relative to control, which was set to 100%. (A**)** *SRA* silencing; (B) *WRB* silencing; (C) *HSND2* silencing; (D) simultaneous *HSND2* plus *WRB* silencing. Notably, the same blot was probed with antibodies against hSnd2, Wrb, and ß-actin respectively, and therefore the same loading control is shown twice.

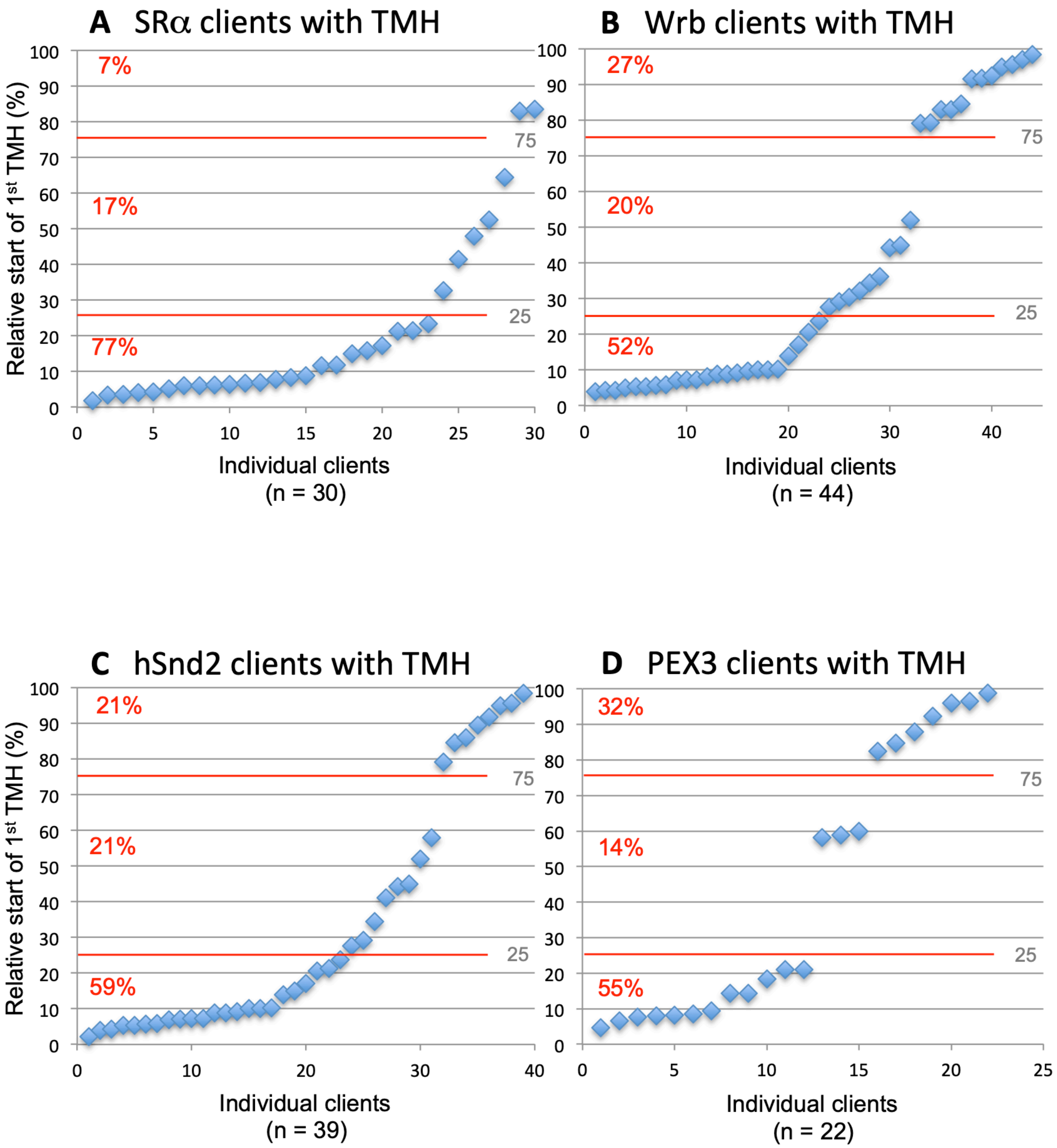
A SRα clients with TMH
B Wrb clients with TMH
C hSnd2 clients with TMH
D PEX3 clients with TMH
Relative start of 1st TMH (%)
Individual clients
(n = 30)
(n = 44)
(n = 39)
(n = 22)
7%
17%
77%
27%
20%
52%
21%
21%
59%
32%
14%
55%
75
25

**Figure S2. Distinguishing Features of SR$\alpha$, Wrb, and hSnd2 Clients with TMH. Related to Figure 2.**

(A-D) TMH containing clients were plotted against the location of their first TMH, i.e. position of central amino acid residue of TMH in % of client amino acid residues. The numbers are shown in Tables S5 and S6.

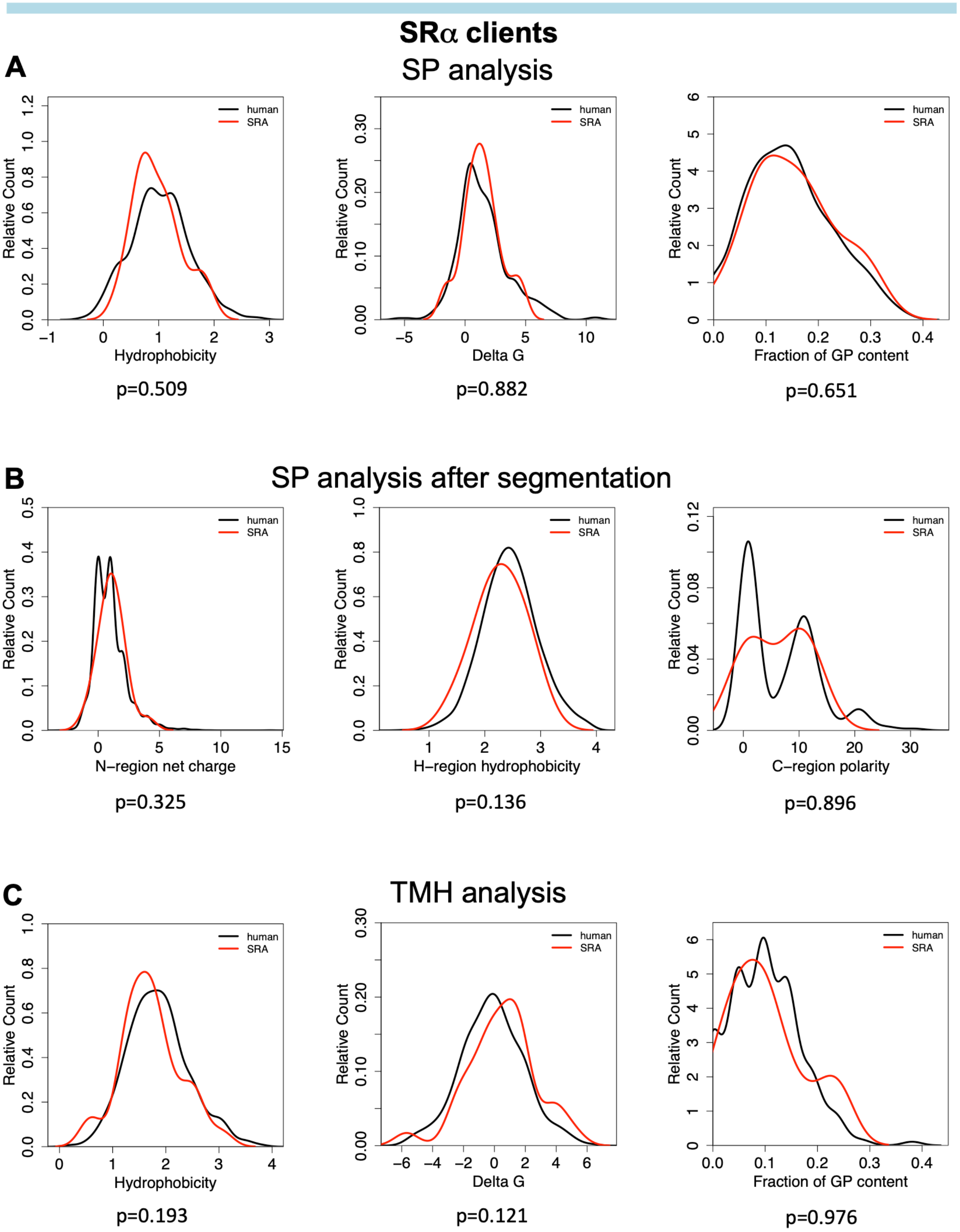
SRα clients
A
SP analysis
Relative Count
human
SRA
Hydrophobicity
p=0.509
Delta G
p=0.882
Fraction of GP content
p=0.651
B
SP analysis after segmentation
N–region net charge
p=0.325
H–region hydrophobicity
p=0.136
C–region polarity
p=0.896
C
TMH analysis
Hydrophobicity
p=0.193
Delta G
p=0.121
Fraction of GP content
p=0.976

**Figure S3. Physicochemical properties of SP, SP-regions and TMH of SR$\alpha$ clients. Related to Figure 2.** (A,C) Using custom scripts, we computed the hydrophobicity and glycine/proline (GP) content of the sequences of SP (A) or TMH (C). The hydrophobicity score of a peptide was calculated as the averaged hydrophobicity of its amino acids according to the well-known Kyte-Doolittle propensity scale as described (Nguyen et al., 2018). GP content was calculated as the total fraction of glycine and proline in the respective sequence as described (Nguyen et al., 2018). We determined the $\Delta G_{app}$ values of SP and TMH with the $\Delta G_{app}$ predictor for transmembrane helix insertion (http://dgpred.cbr.su.se) and plotted all these values against the relative counts. (B) The properties of N-, H- and C-regions of SP of clients were also analyzed after their segmentation by the Phobius (http://phobius.sbc.su.se) prediction tool. Their properties include total net charge of N-region, hydrophobicity of H-region, and the polarity of C-region. Polarity was calculated as the averaged polarity of its amino acids according to the polarity propensity scale as described (Schorr et al., 2020). Likewise, hydrophobicity was calculated using the Kyte-Doolittle propensity scale. We also used custom scripts to extract all SP annotations for human proteins from UniProtKB entries and applied the same calculations for all these human SPs (human).

### Wrb clients

A

#### SP analysis

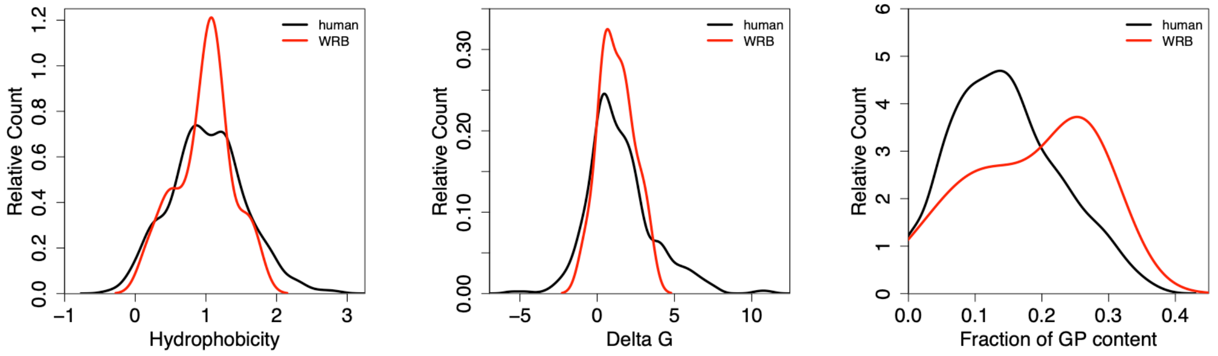

p=0.783 p=0.998 p=0.113

B

#### SP analysis after segmentation

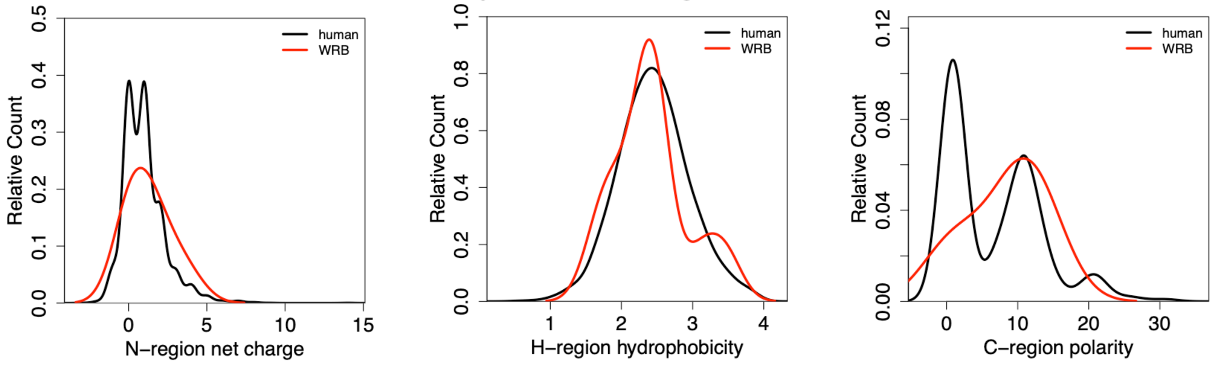

p=0.454 p=0.639 p=0.230

C

#### TMH analysis

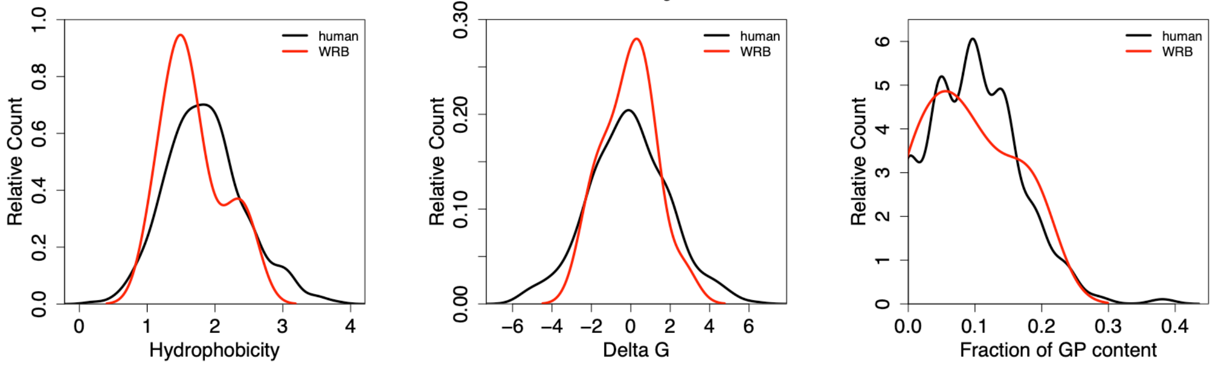

p=0.193 p=0.916 p=0.621

**Figure S4. Physicochemical properties of SP, SP-regions and TMH of Wrb clients. Related to Figure 2.** (A,C) Using custom scripts, we computed the hydrophobicity and glycine/proline (GP) content of the sequences of SP (A) or TMH (C). The hydrophobicity score of a peptide was calculated as the averaged hydrophobicity of its amino acids according to the well-known Kyte-Doolittle propensity scale as described (Nguyen et al., 2018). GP content was calculated as the total fraction of glycine and proline in the respective sequence as described (Nguyen et al., 2018). We determined the $\Delta G_{app}$ values of SP and TMH with the $\Delta G_{app}$ predictor for transmembrane helix insertion (http://dgpred.cbr.su.se) and plotted all these values against the relative counts. (B) The properties of N-, H- and C-regions of SP of clients were also analyzed after their segmentation by the Phobius (http://phobius.sbc.su.se) prediction tool. Their properties include total net charge of N-region, hydrophobicity of H-region, and the polarity of C-region. Polarity was calculated as the averaged polarity of its amino acids according to the polarity propensity scale as described (Schorr et al., 2020). Likewise, hydrophobicity was calculated using the Kyte-Doolittle propensity scale. We also used custom scripts to extract all SP annotations for human proteins from UniProtKB entries and applied the same calculations for all these human SPs (human).

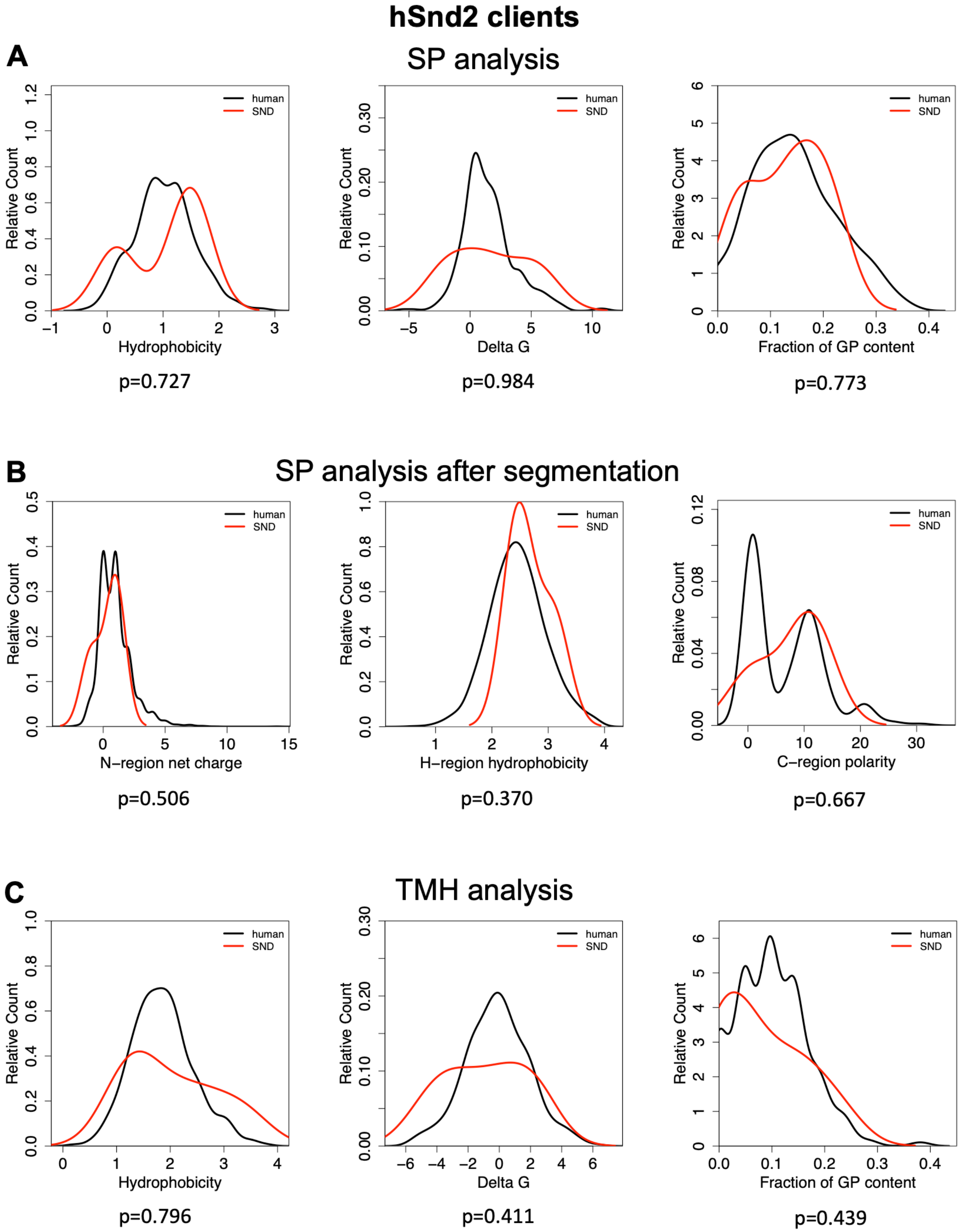
hSnd2 clients
A
SP analysis
Relative Count
Hydrophobicity
human
SND
p=0.727
Delta G
p=0.984
Fraction of GP content
p=0.773
B
SP analysis after segmentation
N–region net charge
p=0.506
H–region hydrophobicity
p=0.370
C–region polarity
p=0.667
C
TMH analysis
Hydrophobicity
p=0.796
Delta G
p=0.411
Fraction of GP content
p=0.439

**Figure S5. Physicochemical properties of SP, SP-regions and TMH of hSnd2 clients. Related to Figure 2.** (A,C) Using custom scripts, we computed the hydrophobicity and glycine/proline (GP) content of the sequences of SP (A) or TMH (C). The hydrophobicity score of a peptide was calculated as the averaged hydrophobicity of its amino acids according to the well-known Kyte-Doolittle propensity scale as described (Nguyen et al., 2018). GP content was calculated as the total fraction of glycine and proline in the respective sequence as described (Nguyen et al., 2018). We determined the $\Delta G_{app}$ values of SP and TMH with the $\Delta G_{app}$ predictor for transmembrane helix insertion (http://dgpred.cbr.su.se) and plotted all these values against the relative counts. (B) The properties of N-, H- and C-regions of SP of clients were also analyzed after their segmentation by the Phobius (http://phobius.sbc.su.se) prediction tool. Their properties include total net charge of N-region, hydrophobicity of H-region, and the polarity of C-region. Polarity was calculated as the averaged polarity of its amino acids according to the polarity propensity scale as described (Schorr et al., 2020). Likewise, hydrophobicity was calculated using the Kyte-Doolittle propensity scale. We also used custom scripts to extract all SP annotations for human proteins from UniProtKB entries and applied the same calculations for all these human SPs (human).

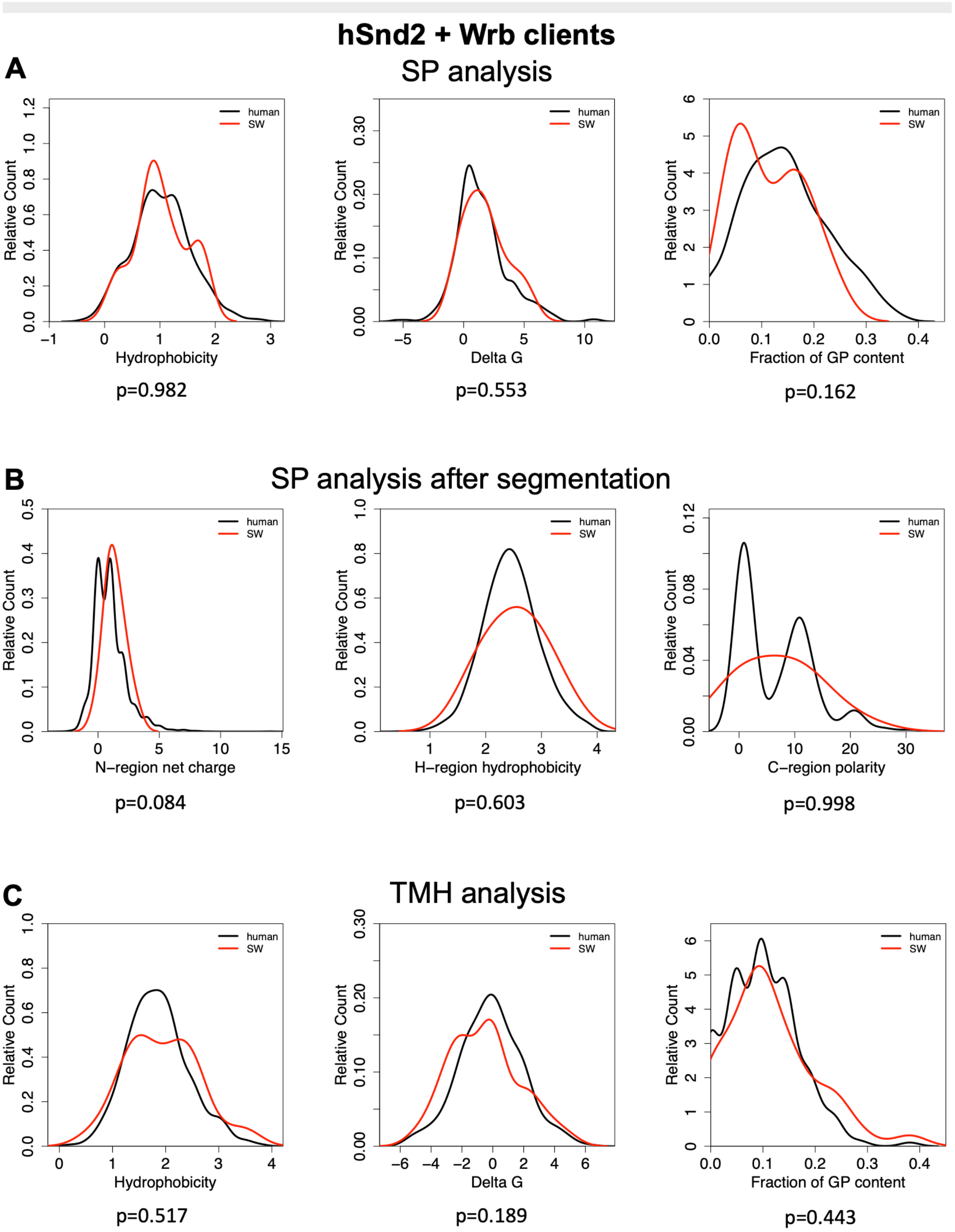
hSnd2 + Wrb clients
A
SP analysis
Relative Count
Hydrophobicity
p=0.982
Delta G
p=0.553
Fraction of GP content
p=0.162
human
SW
B
SP analysis after segmentation
N–region net charge
p=0.084
H–region hydrophobicity
p=0.603
C–region polarity
p=0.998
C
TMH analysis
Hydrophobicity
p=0.517
Delta G
p=0.189
Fraction of GP content
p=0.443

**Figure S6. Physicochemical properties of SP, SP-regions and TMH of hSnd2 plus Wrb clients. Related to Figure 2.** (A,C) Using custom scripts, we computed the hydrophobicity and glycine/proline (GP) content of the sequences of SP (A) or TMH (C). The hydrophobicity score of a peptide was calculated as the averaged hydrophobicity of its amino acids according to the well-known Kyte-Doolittle propensity scale as described (Nguyen et al., 2018). GP content was calculated as the total fraction of glycine and proline in the respective sequence as described (Nguyen et al., 2018). We determined the $\Delta G_{app}$ values of SP and TMH with the $\Delta G_{app}$ predictor for transmembrane helix insertion (http://dgpred.cbr.su.se) and plotted all these values against the relative counts. (B) The properties of N-, H- and C-regions of SP of clients were also analyzed after their segmentation by the Phobius (http://phobius.sbc.su.se) prediction tool. Their properties include total net charge of N-region, hydrophobicity of H-region, and the polarity of C-region. Polarity was calculated as the averaged polarity of its amino acids according to the polarity propensity scale as described (Schorr et al., 2020). Likewise, hydrophobicity was calculated using the Kyte-Doolittle propensity scale. We also used custom scripts to extract all SP annotations for human proteins from UniProtKB entries and applied the same calculations for all these human SPs (human).

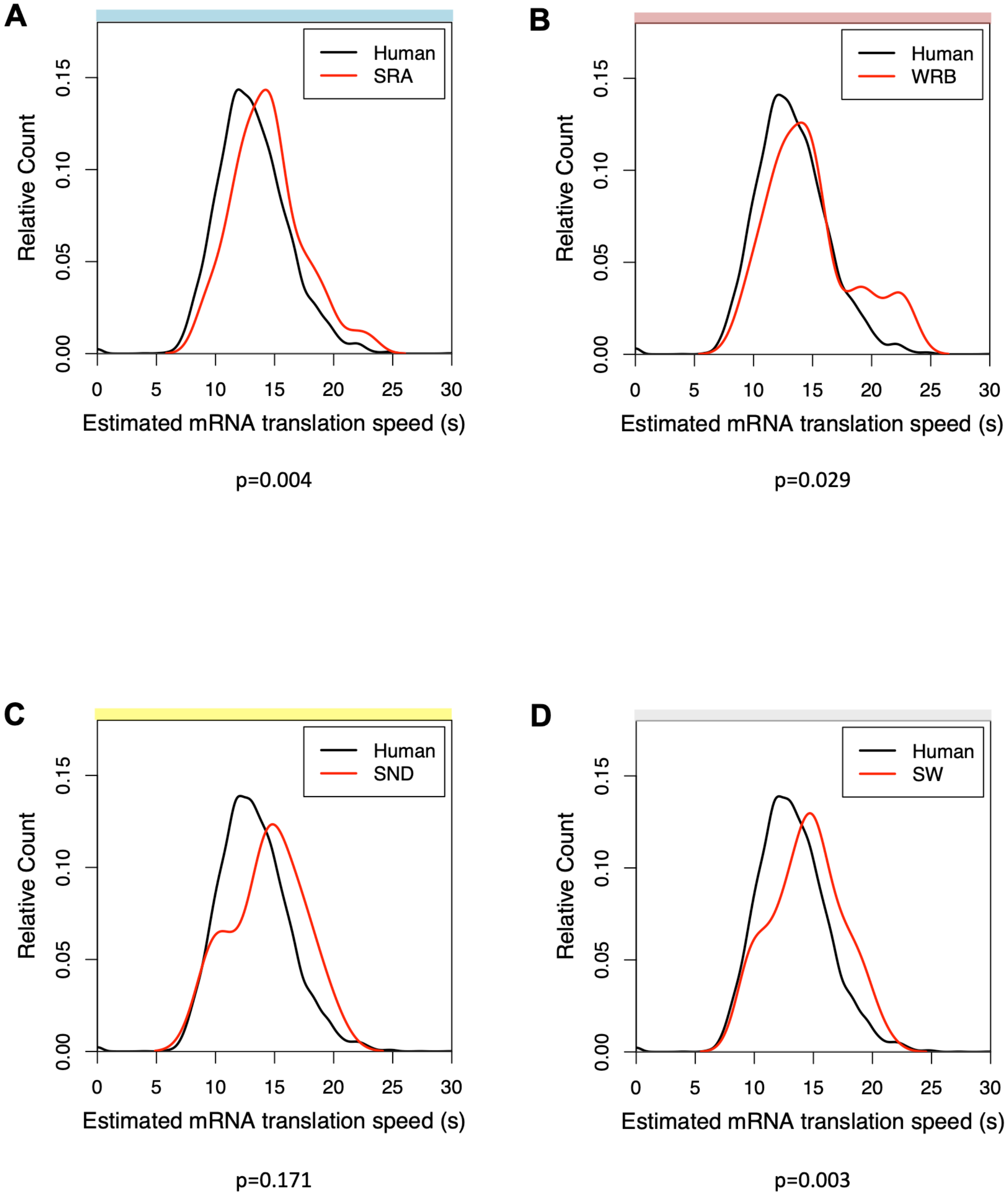
A
B
C
D
Human
SRA
WRB
SND
SW
Relative Count
0.00
0.05
0.10
0.15
0
5
10
15
20
25
30
Estimated mRNA translation speed (s)
p=0.004
p=0.029
p=0.171
p=0.003

**Figure S7. Translation rates of mRNAs coding for clients of SRα, Wrb and hSnd2. Related to Figure 2.** (A-D) The full dataset of human mRNA coding sequences (CDS) was downloaded from www.ensemble.org. It contains 110788 CDS records. To avoid a bias in the comparison between the CDS of clients against the full human CDS (background dataset), we only used the CDS transcripts for the proteins identified in the MS data. For this, we translated the full CDS dataset into protein sequences and compared the resulting proteins with the MS data. 4660 (out of 4856) proteins could be matched in this way. The goal of this analysis is to compare the (estimated) translation rates of the mRNAs negatively affected by knock-down which contain either SP or TMH to the estimated translation rates of the mRNAs in the full MS dataset. As we are only interested in the N-terminal part of the sequence, and to ensure comparability of the results independent of the length of the actual mRNA sequences, we always used the first 240 nucleotides (which is equivalent to 80 amino acids) of the CDS for the subsequent analysis. The translation rates were computed based on the codon-specific elongation rates as provided by Trösenmeier et al., 2019. The rates determined there describe the specific elongation speed for each codon during the translation process. Here, the rates were inverted and summed up, resulting in the (estimated) translation speed for the N-terminal 240 nt of each sequence (240 nt. ~ 80 codons). The two distributions were compared using the nonparametric Wilcoxon test.

**A**

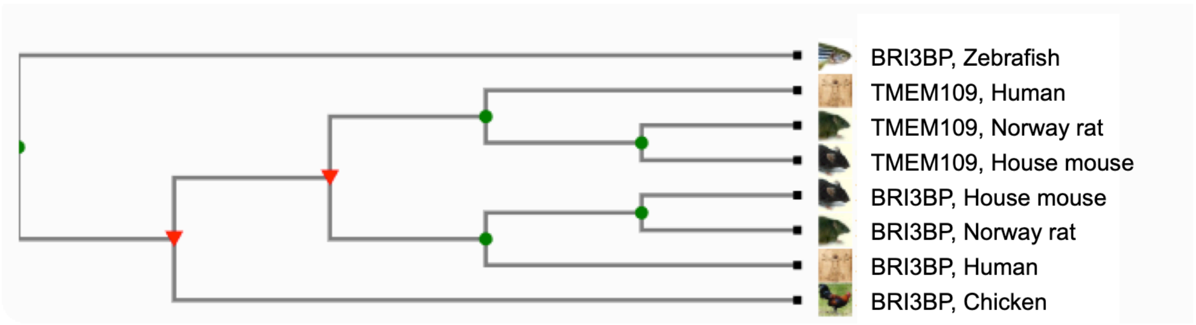

**B**

```
      Signal peptide
1     MAASSISSPWGKHVFKAILMVLVALILLHSALAQSRRDFAPPGQQKREAP          50
      : :     :         :  :  : ::    ::
1     MGARASGGPLARAGLLLLLLLLLLLGLLAPG-AQGARGRGGAEKNSYRRT          49
      Signal peptide
                                       TMD1
51    VDVLTQIGRSVRGTLDAWIGPETMHLVSESSSQVLWAISSAISVAFF--A          100
      :    :   :  :               :              :  :
50    VNTFSQSVSSLFGEDNVRAAQKFLARLTERF---------VLGVDMFVET          90

                                              TMD2
101   LSGIAAQLLNALGLAGDYLAQGLK---LSPGQVQTFLLWGAGALVVYWLLSLLLG     150
             ::  :::    : :       :       ::     :  :: ::: ::
101   LWKVWTELLDVLGLDVSNLSQYFSPASVSSSPARALLLVGVV-LLAYWFLSLTLG     145
                                        TMD1
                                             TMD3
151   LVLALL----GRILWGLKLVIFLAGFVALMRSVPDPSTRALLLLALLI-LYALLS     200
           :    ::  :    :      :              : :      :
151   FTFSVLHVVFGRFFWIVRVVLFSMSCVYILHKYEGEPENAVLPLCFVVAVYFMTG     199
       TMD2                                   TMD3
                        Putative coiled-coil
201   RLT---------GSRASGAQLEAKVRGLERQVEELRWRQRRAAKGARSVEEE        243
                   :  :    : :   :: ::  :  :  :
201   PMGFYWRSSPSGPSNPSNPSVEEKLEHLEKQVRLLNIRLNRVLESLDRSKDK        251
                          Putative coiled-coil
```

**Figure S8. TMEM109 and BRI3BP are part of a protein family. Related to Figure 5.** (A) The phylogenetic tree was generated according to Ruan et al., 2008 and Guindon et al., 2010, respectively, by the TreeFam option of http://pfam.xfam.org. (B) The sequences of human TMEM109 (Q9BVC6) and BRI3BP (Q8WY22) were extracted from UniprotKB. Sequence alignment was performed with the ClustalV option of the MegAlign tool within the DNASTAR sofware package (Lasergene 12). Signal peptides, transmembrane domains (TMD) and coiled coil domains are indicated as determined by UniProtKB, https://services.healthtech.dtu.dk/service.php?SignalP and http://gpcr.biocomp.unibo.it/cgi/predictors/cc/pred_cchmm.cgi, respectively.

**Table S1, related to Figure 1A.**

File Name: Table S1_sra_all.xlsx

Complete list of genes corresponding to proteins quantified after *SRA* silencing in HeLa cells. Gene names, protein accession numbers (ID), log2 fold changes resulting from siRNA-mediated SRα depletion, and -log10 p values are indicated. Minus sign in front of fold change denotes negatively affected proteins. The number of listed proteins differs from the total number of quantified proteins because some proteins were quantified in less than two of the triplicates. The original Orbitrap data for all quantified proteins are deposited at Proteome Exchange: http://www.proteomexchange.org.

**Table S2, related to Figure 1A.**

File Name: Table S2_sra_full_lo.xlsx

Proteins that were negatively affected by *SRA* silencing in HeLa cells, i.e. putative SRα clients. Gene names, protein accession numbers (ID), and log2 fold changes resulting from siRNA-mediated SRα depletion are presented together with full protein names and Gene Ontology (GO) annotations for subcellular location(s), presence of N-terminal signal peptide (SP) or most N-terminal transmembrane helix (TMH), number of N-glycosylation sites (Glycosylation sites), number of transmembrane domains (TMD), amino acid sequences of SP or TMH (in single letter code), position of TMH, all as extracted from UniProtKB entries using custom scripts. Proteins are listed according to decreasing negative effects of SRα depletion.

**Table S3, related to Figure 1A.**

File Name: Table S3_sra_full_up.xlsx

Proteins that were positively affected by *SRA* silencing in HeLa cells. Gene names, protein accession numbers, and log2 fold changes resulting from SRα depletion are presented together with Gene Ontology (GO) annotations for subcellular location(s), presence of N-terminal signal peptide (SP) or N-terminal transmembrane helix (TMH), number of N-glycosylation sites (Glycosylation sites), amino acid sequences of SP or TMH (in single letter code), all as extracted from UniProtKB entries using custom scripts. Proteins are listed according to decreasing positive effects of SRα depletion.

**Table S4, related to Figure 2. Summary of clients as determined by MS and differential protein abundance analysis.**

| Clients of | Clients with | | | Clients with | | | | |
|---|---|---|---|---|---|---|---|---|
| | SP | SP/I | SP/multi | TMH | TMH/II or III | TMH/multi | HP | TA |
| **SRα** | **24** | **14** | **-** | **30** | **10** | **17** | **3** | **-** |
| Total | 54 | | | | | | | |
| % | 44.4 SP | | | 55.6 TMH | | | | |
| % | 81.5 MP | | | | | | | |
| % of TMH | | | | | 33.3 | 56.7 | 10 | 0 |
| % | 18.5sol | 25.9 | | | 18.5 | 31.5 | 5.6 | 0 |
| **Wrb** | **13** | **6** | **-** | **14** | **2** | **8** | **1** | **3** |
| Total | 27 | | | | | | | |
| % | 48.1 SP | | | 51.9 TMH | | | | |
| % | 74.1 MP | | | | | | | |
| % of TMH | | | | | 14.3 | 57.1 | 7.1 | 21.4 |
| % | | | | | 7.4 | 29.6 | 3.7 | 10.1 |
| **hSnd2** | **3** | **3** | **-** | **9** | **2** | **5** | **-** | **2** |
| Total | 12 | | | | | | | |
| % | 25 SP | | | 75 TMH | | | | |
| % | 100 MP | | | | | | | |
| % of TMH | | | | | 22.2 | 55.6 | 0 | 22.2 |
| % | | | | | 16.7 | 41.7 | 0 | 16,7 |
| **hSnd2+Wrb** | **13** | **5** | **2** | **30** | **3** | **21** | **1** | **5** |
| Total | 43 | | | | | | | |
| % | 30.2 SP | | | 69.8 TMH | | | | |
| % | 86 MP | | | | | | | |
| % of TMH | | | | | 10 | 70 | 3.3 | 16.7 |
| % | | | | | 7 | 48.8 | 2.3 | 11.6 |
| **Wrb** sum | **26** | **11** | **2** | **44** | **5** | **29** | **2** | **8** |
| Total | 70 | | | | | | | |
| % | 37.1 SP | | | 62.9 TMH | | | | |
| % | 81.4 MP | | | | | | | |
| % of TMH | | | | | 11.4 | 65.9 | 4.5 | 18.2 |
| % | 18.6sol | 15.7 | 2.9 | | 7.1 | 41.4 | 2.9 | 11.4 |
| **hSnd2** sum | **16** | **8** | **2** | **39** | **5** | **26** | **1** | **7** |
| Total | 55 | | | | | | | |
| % | 29,1 SP | | | 70.9 TMH | | | | |
| % | 89.1 MP | | | | | | | |
| % of TMH | | | | | 12.8 | 66.7 | 2.6 | 17.9 |
| % | 10.9sol | 14.5 | 3.6 | | 9.1 | 47.3 | 1.8 | 12.7 |
| **PEX3** | 1 | - | 1 | 2 | 1 | 1 | - | - |
| Zellweger | 27 | 5 | - | 20 | 8 | 6 | 2 | 4 |
| | **28** | **5** | **1** | **22** | **9** | **7** | **2** | **4** |
| Total | 50 | | | | | | | |
| % | 56 SP | | | 44 TMH | | | | |
| % | 56 MP | | | | | | | |
| % of TMH | | | | | 40.9 | 31.8 | 9.1 | 18.2 |
| % | 44sol | 10 | 2 | | 18 | 14 | 4 | 8 |

I, II, III, membrane protein type; HP, hairpin; MP, membrane protein; multi, multispanning membrane protein; sol, soluble; SP, signal peptide; sum, sum of clients as determined in single depletion and double depletion (hSnd2+Wrb); TA, tail anchor; TMH, most N-terminal transmembrane domain; Zellweger, data from PEX3 deficient patient fibroblasts (Zimmermann et al., 2021).

#### Table S5, related to Figure 2. Characteristics of clients of SRα, Wrb, and hSnd2 with TMH.

| | TMDs | Type | ΔGapp | Sequence | 1st TMH | Size | % | N-glyco |
|---|---|---|---|---|---|---|---|---|
| **SRα** | | | | | | | | |
| ABCC4 | multi | | 5.118 | LVLGIFTLIEESAKVIQPIFL | 93-113 | 1325 | 7.8 | + |
| ANO10 | multi | | 1.133 | IALYFGFLEYFTFALIPMAVI | 208-228 | 660 | 32.7 | |
| ASPH | 1 | II | -1.493 | FFTWFMVIALLGVWTSVAVVW | 54-74 | 758 | 8.4 | + |
| ATL2 | 1 | HP | -0.074 | TLFAVMFAMYIISGLTGFIGL | 477-497 | 583 | 83.5 | + |
| ATP2B1 | multi | | 4.019 | FLQLVWEALQDVTLIILEIAA | 98-118 | 1220 | 8.9 | |
| ATP13A1 | multi | | 1.928 | VLPFAGLLYPAWLGAAAAGCW | 67-87 | 1204 | 6.4 | + |
| B3GALTL | 1 | II | -0.935 | WWLLAPPALLALLTCSLAFGL | 7-27 | 498 | 3.4 | + |
| BST2 | 1 | II | -5.685 | KLLLGIGILVLLIIVILGVPLIIFTIKA | 21-48 | 180 | 17.2 | + |
| CAV1 | 1 | HP | -0.726 | ALFGIPMALIWGIYFAILSFL | 105-125 | 178 | 64.4 | |
| CEPT1 | multi | | -0.476 | LITIIGLSINICTTILLVFYC | 87-107 | 416 | 23.3 | + |
| DEGS1 | multi | | -0.478 | PNLIWIIIMMVLTQLGAFYIV | 41-61 | 323 | 15.8 | |
| ERGIC2 | multi | | 1.141 | GTVSLIAFTTMALLTIMEFSV | 34-54 | 377 | 11.7 | |
| ERLIN2 | 1 | II | 1.947 | LGAVVAVASSFFCASLFSAVH | 4-24 | 339 | 4.1 | + |
| IKBIP | 1 | | 0.469 | CLSLLSLGTCLGLAWFV | 46-62 | 377 | 14.9 | + |
| ITPR3 | multi | | 0.138 | LWGSISFNLAVFINIIIAFFY | 2203-2223 | 2671 | 82.9 | + |
| LNPEP | 1 | II | -2.599 | MVVCAFVIVVAVSVIMVIYLL | 111-131 | 1025 | 11.8 | |
| PDE3A | multi | | -0.598 | LSSALCAGSLSFLLALLVRLV | 61-81 | 1141 | 6.2 | |
| PEX3 | 1 | | 3.594 | CIFLGTVLGGVYILGKYGQKK | 16-36 | 373 | 7 | |
| REEP3 | multi | HP | 0.697 | MVSWMISRAVVLVFGMLYPAY | 1-21 | 255 | 4.3 | |
| SLC16A3 | multi | | -0.663 | GGWGWAVLFGCFVITGFSYAF | 18-38 | 465 | 6 | |
| SLC35B2 | multi | | 3.884 | WWAVVVLAAFPSLGAGGETPE | 5-25 | 432 | 3.5 | |
| SPTLC1 | 1 | | 1.444 | ALYEAPAYHLILEGILILWII | 16-36 | 513 | 5.1 | |
| SUN1 | 1 | II | -1.860 | ICKFLVLLIPLFLLLAGLSL | 316-335 | 785 | 41.5 | |
| TMEM41B | multi | | -1.951 | MSLLILVSIFLSAAFVMFLVY | 52-72 | 291 | 21.3 | |
| TMEM209 | multi | | 1.532 | VVLAWGLLNVSMAGMIYTEMT | 28-48 | 561 | 6.8 | + |
| TMTC3 | multi | | 1.675 | ITLIVGVVTACYWNSLFCGFV | 9-29 | 914 | 1.8 | + |
| TOR1AIP2 | 1 | | -0.371 | FWSYGPVILVVLVVAVVASSV | 215-235 | 470 | 47.9 | + |
| TVP23B | multi | | 1.886 | PVASFFHLFFRVSAIIVYLL | 34-53 | 205 | 21.5 | |
| YIPF5 | multi | | 0.365 | TDLAGPMVFCLAFGATLLLAG | 125-145 | 257 | 52.5 | |
| ZMPSTE24 | multi | | 1.532 | IFGAVLLFSWTVYLWETFLAQ | 19-39 | 475 | 6.1 | |
| **Wrb** | | | | | | | | |
| EXT2 | 1 | II | -1.428 | YITLFSIVLLGLIATGMFQFW | 26-46 | 718 | 5 | |
| FAR1 | 1 | TA | -0.255 | IRYGFNTILVILIWRIF | 466-483 | 515 | 92.4 | |
| GOLGA5 | 1 | TA | -2.261 | VFVIIYMALLHLWVMIVLLTYTP | 699-719 | 731 | 97 | |
| ITPR3 | multi | | 0.138 | LWGSISFNLAVFINIIIAFFY | 2203-2223 | 2671 | 82.9 | |
| LBR | multi | | -2.378 | VPGVFLIMFGLPVFLFLLLLM | 212-232 | 615 | 36.1 | |
| MARCH1 | multi | | 0.107 | IFCSVTFHVIAITCVVWSLYV | 155-175 | 545 | 30.3 | |
| MFSD7 | multi | | 0.335 | WVFLLAISLLNCSNATLWLSF | 30-50 | 559 | 7.2 | |
| MFSD10 | multi | | -1.459 | VVFLGLLLDLLAFTLLLPLLP | 27-47 | 455 | 8.1 | |
| NEU1 | 1 | | -1.467 | LGFWGGCRVWVFAAIFLLLSLAA | 20-41 | 415 | 7.2 | + |
| PTDSS1 | multi | | 0.972 | FFYRPHTITLLSFTIVSLMYF | 36-56 | 473 | 9.7 | |
| REEP3 | multi | HP | 0.697 | MVSWMISRAVVLVFGMLYPAY | 1-21 | 255 | 4.3 | |
| SGPP1 | multi | | 2.661 | FCFGTELGNELFYILFFPFWI | 132-152 | 441 | 32.2 | |
| SPCS2 | multi | | 1.255 | ICTISCFFAIVALIWDYMHPF | 87-107 | 226 | 82.9 | |
| UBE2J2 | 1 | TA | -1.019 | GLLGGALANLFVIVGFAAFAYTW | 227-247 | 259 | 91.5 | |
| **hSnd2** | | | | | | | | |
| MYO9A | 1 | | 1.571 | IYTYVGSILIVINPFKFLPIY | 175-195 | 2548 | 7.3 | |
| PTDSS2 | multi | | -0.192 | AHTLTVLFILTCTLGYVTLLE | 63-83 | 487 | 15 | + |
| PTGIS | 1 | | -3.435 | MAWAALLGLLAALLLLLLLS | 1-20 | 500 | 2.2 | |
| SLC4A2 | multi | | -0.458 | CLAAVIFIYFAALSPAITFGGLLG | 708-731 | 1241 | 57.9 | + |
| SLC7A2 | multi | | 2.693 | DLIALGVGSTLGAGVYVLAGEV | 38-59 | 658 | 7.3 | + |
| TMEM41B | multi | | -1.951 | MSLLILVSIFLSAAFVMFLVY | 52-72 | 291 | 21.3 | |
| TRPM7 | multi | | 1.975 | NSWYKVILSILVPPAILLLEY | 756-776 | 1865 | 41.1 | |
| VAMP4 | 1 | TA | -3.47 | IKAIMALVAAILLLVIIILIV | 116-136 | 141 | 89.4 | |
| VAMP8 | 1 | TA | -4.429 | MVLICVIVFIIILFIVLFAT | 76-96 | 100 | 86 | |

1st TMH, most N-terminal transmembrane domain; TMD, number of transmembrane domains (including TMH); 1, single spanning; multi, multispanning; Type, membrane protein type; HP, hairpin; TA, tail anchor; ΔGapp, apparent delta G of TMH; Sequence, primary structure of TMH; amino acid residues of TMH; Size, number of amino acid residues of client; %, distribution of TMH in client (i.e. position of central amino acid residue of TMH in % of client); N-glyco, N-glycosylation.

**Table S6, related to Figure 2. Characteristics of clients of Wrb plus hSnd2 and PEX3 with TMH.**

| | TMDs | Type | ΔGapp | Sequence | TMH | Size | % | N-glyco |
|---|---|---|---|---|---|---|---|---|
| **hSnd2+Wrb** | | | | | | | | |
| AGPAT5 | multi | | 1.565 | LLPSVVLLGTAPTYVLAWGVW | 15-35 | 364 | 6.9 | |
| ATG9A | multi | | -0.147 | IFELMQFLFVVAFTTFLVSCV | 67-87 | 839 | 9.2 | + |
| ATP2C1 | multi | | 4.913 | LWKKYISQFKNPLIMLLLASA | 71-91 | 919 | 8.8 | |
| ATP12A | multi | | 3.561 | EIVKFLKQMVGGFSILLWVGA | 102-123 | 1093 | 10.3 | |
| BCAP29 | multi | | -0.863 | AVATFLYAEIGLILIFCLPFI | 7-27 | 241 | 7.1 | |
| C4orf3 | 1 | TA | -2.704 | SYWLDLWLFILFDVVVFLFVYFL | 45-65 | 65 | 84.6 | |
| CXCR4 | multi | | -0.37 | IFLPTIYSIIFLTGIVGNGLVILVM | 39-63 | 352 | 13.9 | + |
| EMD | 1 | TA | -2.850 | VPLWGQLLLFLVFVIVLFFIY | 223-243 | 254 | 91.7 | |
| GDPD4 | multi | | 0.185 | WVTFLGTGYWFFWSIFILSLA | 18-38 | 520 | 5.4 | + |
| JPH1 | 1 | TA | -1.838 | IMIVLVMLLNIGLAILFVHFL | 640-660 | 661 | 98.3 | |
| LEMD2 | multi | | -3.464 | LLLWASLGLLLVFLGILWVKM | 213-233 | 503 | 44.3 | |
| MBOAT7 | multi | | 2.483 | LVVLLISIPIGFLFKKAGPGL | 9-29 | 472 | 4 | + |
| MXRA7 | 1 | | 2.128 | LLAALPALATALALLLAWLLV | 7-27 | 170 | 10 | |
| PLD3 | 1 | II | -1.979 | VLLVLILAVVGFGALMTQLFL | 39-59 | 490 | 10 | + |
| POMK | 1 | II | -0.095 | VGLLLIMALMNTLLYLCLDHFFI | 21-43 | 350 | 8.9 | + |
| PRAF2 | multi | | 0.209 | LYYQTNYLLCFGIGLALAGYV | 42-62 | 178 | 29.2 | |
| RHBDD2 | multi | | 2.164 | WCLCPEVPSATFFTALLSLLV | 11-31 | 364 | 5.8 | |
| REEP5 | multi | HP | -1.133 | SFIALGVIGLVALYLVFGYGA | 35-55 | 189 | 23.8 | |
| SEC62 | multi | | -1.555 | FVMGLILVIAVIAATLFPLWP | 197-217 | 399 | 51.9 | |
| SLC9A6 | multi | | 3.829 | LWLLLAVGVFDWAGASDGGGG | 28-48 | 679 | 5.6 | + |
| SLC16A7 | multi | | -0.035 | GGWGWIVVGAAFISIGFSYAF | 16-36 | 478 | 5.4 | |
| SLC39A7 | multi | | 0.193 | WVAVGLLTWATLGLLVAGLGG | 10-30 | 469 | 4.3 | |
| SOAT1 | multi | | -2.080 | IYHMFIALLILFILSTLVV | 141-159 | 550 | 27.5 | |
| STEAP4 | multi | | -0.321 | LFPMWRFPFYLSAVLCVFLFF | 196-216 | 459 | 44.9 | |
| STX2 | 1 | TA | -2.654 | WIIIAVSVVLVAIIALIIGLSVGK | 265-288 | 288 | 95.5 | |
| STX3 | 1 | TA | -4.337 | LIIIIVLVVVLLGILALIIGLSV | 264-284 | 289 | 94.8 | |
| STX17 | multi | | -2.116 | LAALPVAGALIGGMVGGPIGL | 229-249 | 302 | 79.1 | |
| TMEM33 | multi | | 0.388 | LFTVYCSALFVLPLLGLHEAA | 32-52 | 247 | 17 | + |
| TMEM38B | multi | | 0.584 | SWFTAMLHCFGGGILSCLLLA | 50-70 | 291 | 20.6 | |
| TMEM181 | multi | | -2.195 | HFVLVFVVFFICFGLTIFVGI | 153-173 | 475 | 34.4 | |
| **PEX3** | | | | | | | | |
| ABCD3 | multi | | 1.380 | GYLVLIAVMLVSRTYCDVWMI | 84-104 | 659 | 14.3 | + |
| ACBD5 | 1 | | -0.121 | GVLTFAIIWPFIAQWLVYLYY | 497-517 | 525 | 96.6 | |
| AIFM2 | 1 | | 2.886 | VESGALHVVIVGGGFGGIAAA | 7-27 | 373 | 4.6 | |
| ATL1 | 1 | HP | -0.573 | TLFVVIFITYVIAGVTGFIGL | 450-470 | 558 | 82.4 | |
| CCDC136 | 1 | TA | -2.371 | IFSLPLVGLVVISALLWCWWA | 1130-1150 | 1154 | 98.8 | |
| COLEC12 | 1 | II | -2.162 | FSIILLYILCALLTITVAILG | 38-58 | 742 | 6.5 | + |
| CYBRD1 | multi | | -2.348 | LLGSALLVGFLSVIFALVWVL | 12-32 | 286 | 7.7 | + |
| DHRS7B | 1 | II | -0.36 | FITSTAILPLLFGCLGVFGLF | 18-38 | 325 | 8.6 | |
| ENPP1 | 1 | II | -0.985 | VLSLVLSVCVLTTILGCIFGL | 77-97 | 925 | 9.4 | + |
| ERMP1 | multi | | 2.156 | AGTGLSEVRAALGLALYLIAL | 64-84 | 904 | 8.2 | + |
| FAR1 | 1 | TA | -0.668 | IRYGFNTILVILIWRIFI | 466-483 | 515 | 92.4 | |
| MAN1A1 | 1 | II | -2.612 | FVLLLVFSAFITLCFGAIFFL | 42-62 | 653 | 8 | + |
| PEX13 | 1 | | 0.039 | AATSAKSWPIFLFFAVILGGPYLIW | 227-251 | 403 | 58.8 | |
| PXMP2 | multi | | 3.630 | LYPVLTKAATSGILSALGNFL | 31-51 | 195 | 21 | |
| RTN3 | multi | HP | -2.436 | LIMLLSLAAFSVISVVSYLILALL | 864-887 | 1032 | 84.7 | |
| SGCD | 1 | II | -3.808 | FFVLLLMILILVNLAMTIWIL | 37-57 | 256 | 18.4 | |
| STX6 | 1 | TA | -5.096 | WCAIAILFAVLLVVLILFLVL | 235-255 | 255 | 96.1 | + |
| TMEM192 | multi | | -1.154 | TVIIVNLLWFIHLVFVVLAFL | 47-67 | 271 | 21 | |
| TMEM237 | multi | | 1.917 | MIGLFSHGFLAGCAVWNIVVI | 227-247 | 408 | 58.1 | |
| TMUB2 | multi | | -2.382 | VMVVAGVVVLILALVLAWLST | 36-56 | 321 | 14.3 | |
| TOR1AIP1 | 1 | | 0.705 | WLLPLIAALASGSFWFF | 339-355 | 583 | 59.9 | |
| VAMP3 | 1 | TA | -4.236 | MWAIGITVLVIFIIIIIVWVV | 78-98 | 100 | 88 | |

1st TMH, most N-terminal transmembrane domain; TMD, number of transmembrane domains (including TMH); 1, single spanning; multi, multispanning; Type, membrane protein type; HP, hairpin; TA, tail anchor; ΔGapp, apparent delta G of TMH; Sequence, primary structure of TMH; amino acid residues of TMH; Size, number of amino acid residues of client; %, distribution of TMH in client (i.e. position of central amino acid residue of TMH in % of client); N-glyco, N-glycosylation.

**Table S7, related to Figure 1B.**

File Name: Table S4_wrb_all.xlsx

Complete list of genes corresponding to proteins quantified after *WRB* silencing in HeLa cells. Gene names, protein accession numbers (ID), log2 fold changes resulting from siRNA-mediated Wrb depletion, and -log10 p values are indicated. Minus sign in front of fold change denotes negatively affected proteins. The number of listed proteins differs from the total number of quantified proteins because some proteins were quantified in less than two of the triplicates. The original Orbitrap data for all quantified proteins are deposited at Proteome Exchange: http://www.proteomexchange.org.

**Table S8, related to Figure 1B.**

File Name: Table S5_wrb_full_lo.xlsx

Proteins that were negatively affected by *WRB SRA* silencing in HeLa cells, i.e. putative Wrb clients. Gene names, protein accession numbers (ID), and log2 fold changes resulting from siRNA-mediated Wrb depletion are presented together with full protein names and Gene Ontology (GO) annotations for subcellular location(s), presence of N-terminal signal peptide (SP) or most N-terminal transmembrane helix (TMH), number of N-glycosylation sites (Glycosylation sites), number of transmembrane domains (TMD), amino acid sequences of SP or TMH (in single letter code), position of TMH, all as extracted from UniProtKB entries using custom scripts. Proteins are listed according to decreasing negative effects of Wrb depletion.

**Table S9, related to Figure 1B.**

File Name: Table S6_wrb_full_up.xlsx

Proteins that were positively affected by *WRB* silencing in HeLa cells. Gene names, protein accession numbers, and log2 fold changes resulting from Wrb depletion are presented together with Gene Ontology (GO) annotations for subcellular location(s), presence of N-terminal signal peptide (SP) or N-terminal transmembrane helix (TMH), number of N-glycosylation sites (Glycosylation sites), amino acid sequences of SP or TMH (in single letter code), all as extracted from UniProtKB entries using custom scripts. Proteins are listed according to decreasing positive effects of Wrb depletion.

**Table S10, related to Figure 3A.**

File Name: Table S7_snd_all.xlsx

Complete list of genes corresponding to proteins quantified after *hSND2* silencing in HeLa cells. Gene names, protein accession numbers (ID), log2 fold changes resulting from siRNA-mediated hSnd2 depletion, and -log10 p values are indicated. Minus sign in front of fold change denotes negatively affected proteins. The number of listed proteins differs from the total number of quantified proteins because some proteins were quantified in less than two of the triplicates. The original Orbitrap data for all quantified proteins are deposited at Proteome Exchange: http://www.proteomexchange.org.

**Table S11, related to Figure 3A.**

File Name: Table S8_snd_full_lo.xlsx

Proteins that were negatively affected by *hSND2* silencing in HeLa cells, i.e. putative hSND2 clients. Gene names, protein accession numbers (ID), and log2 fold changes resulting from siRNA-mediated hSnd2 depletion are presented together with full protein names and Gene Ontology (GO) annotations for subcellular location(s), presence of N-terminal signal peptide (SP) or most N-terminal transmembrane helix (TMH), number of N-glycosylation sites (Glycosylation sites), number of transmembrane domains (TMD), amino acid sequences of SP or TMH (in single letter code), position of TMH, all as extracted from UniProtKB entries using custom scripts. Proteins are listed according to decreasing negative effects of hSnd2 depletion.

**Table S12, related to Figure 3A.**

File Name: Table S9_snd_full_up.xlsx

Proteins that were positively affected by *hSND2* silencing in HeLa cells. Gene names, protein accession numbers, and log2 fold changes resulting from hSnd2 depletion are presented together with Gene Ontology (GO) annotations for subcellular location(s), presence of N-terminal signal peptide (SP) or N-terminal transmembrane helix (TMH), number of N-glycosylation sites (Glycosylation sites), amino acid sequences of SP or TMH (in single letter code), all as extracted from UniProtKB entries using custom scripts. Proteins are listed according to decreasing positive effects of hSnd2 depletion.

**Table S13, related to Figure 3B.**

File Name: Table S10_sw_all.xlsx

Complete list of genes corresponding to proteins quantified after simultaneous *hSND2* and *WRB* silencing in HeLa cells. Gene names, protein accession numbers (ID), log2 fold changes resulting from siRNA-mediated hSnd2 and Wrb depletion, and -log10 p values are indicated. Minus sign in front of fold change denotes negatively affected proteins. The number of listed proteins differs from the total number of quantified proteins because some proteins were quantified in less than two of the triplicates. The original Orbitrap data for all quantified proteins are deposited at Proteome Exchange: http://www.proteomexchange.org.

**Table S14, related to Figure 3B.**

File Name: Table S11_sw_full_lo.xlsx

Proteins that were negatively affected by simultaneous *hSND2* and *WRB* silencing in HeLa cells, i.e. putative hSnd2 and Wrb clients. Gene names, protein accession numbers (ID), and log2 fold changes resulting from siRNA-mediated hSnd2 and Wrb depletion are presented together with full protein names and Gene Ontology (GO) annotations for subcellular location(s), presence of N-terminal signal peptide (SP) or most N-terminal transmembrane helix (TMH), number of N-glycosylation sites (Glycosylation sites), number of transmembrane domains (TMD), amino acid sequences of SP or TMH (in single letter code), position of TMH, all as extracted from UniProtKB entries using custom scripts. Proteins are listed according to decreasing negative effects of hSnd2 and Wrb depletion.

**Table S15, related to Figure 3B.**

File Name: Table S12_sw_full_up.xlsx

Proteins that were positively affected by simultaneous *hSND2* and *WRB* silencing in HeLa cells. Gene names, protein accession numbers, and log2 fold changes resulting from hSnd2 and Wrb depletion are presented together with Gene Ontology (GO) annotations for subcellular location(s), presence of N-terminal signal peptide (SP) or N-terminal transmembrane helix (TMH), number of N-glycosylation sites (Glycosylation sites), amino acid sequences of SP or TMH (in single letter code), all as extracted from

UniProtKB entries using custom scripts. Proteins are listed according to decreasing positive effects of hSnd2 and Wrb depletion.

**Table S16. Related to Figure 5.**

File Name: Table S16_Overlap siSND, siSND+WRB, CO-IP hSnd2.xlsx

**Table S17, related to STAR Methods.** ProteomeXchange_identifiers

| | | |
|---|---|---|
| Orbi1986 | SRA experiment 1 | PXD008178 |
| Sample 1-3 | scr control siRNA | |
| Sample 4-6 | SRA siRNA #3 | |
| Sample 7-9 | SRA siRNA #6 | |
| Orbi2048 | WRB | PXD008178 |
| Sample 16-18 | scr control siRNA | |
| Sample 19-21 | WRB siRNA #3 | |
| Sample 22-24 | WRB siRNA #4 | |
| Orbi2155 | SRA experiment 2 | PXD012078 |
| Sample 1-3 | scr control siRNA | |
| Sample 10-12 | SRA siRNA #3 | |
| Sample 13-15 | SRA siRNA #6 | |
| Orbi2514 | SND2 & WRB+SND2 experiment 2 | PXD011993 |
| Sample 1-3 | scr control siRNA | |
| Sample 4-6 | SND2 siRNA #2 | |
| Sample 7-9 | SND2 siRNA #3 | |
| Sample 10-12 | WRB siRNA #3 + SND2 siRNA #2 = SW | |
| Sample 13-15 | WRB siRNA #3 + SND2 siRNA #3 = SW | |

The mass spectrometry proteomics data (.raw and .txt files) have been deposited to the ProteomeXchange Consortium via the PRIDE partner repository with the dataset identifiers: PXD008178, PXD011993 and PXD012078 (http://www.proteomexchange.org).

**Table S18, related to STAR Methods. siRNAs used in this study.**

| Name | Target sequence | Source | Concentration (nM) | Time (h) |
|---|---|---|---|---|
| *HSND2*-UTR siRNA #2 | CTCTATAGGGTCGTTGAATAA | Qiagen | 20 | 96 |
| *HSND2* siRNA #3 | AAGGGCAAAGTGGGCACGAGA | Qiagen | 20 | 96 |
| *SRA*-UTR siRNA #3 | CACCAGAGCTTTGCTAATAAT | Qiagen | 15 | 96 |
| *SRA*-UTR siRNA #6 | CAGAGAAATAAGTAATTTATA | Qiagen | 15 | 96 |
| *TMEM109*-UTR #1 | CAGGTTTGATGTGGAATCACA | Qiagen | 20 | 72 |
| *TMEM109-UTR #2* | CACCGCCAGTGTCATACCAAA | Qiagen | 20 | 72 |
| *WRB*-UTR siRNA #3 | TGACACGTATGTACTAGTGAA | Qiagen | 20 | 96 |
| *WRB* siRNA #4 | CACAGTCAACATGATGGACGA | Qiagen | 20 | 96 |

**Table S19, related to STAR Methods. PCR primers used to create plasmids for live cell protein-protein-interaction.**

| Construct | Fwd primer (5´-3´) | Rev primer (5´-3´) |
|---|---|---|
| hSnd2-$C_S$ | TAATACGACTCACTATAGG | TCCCGGTG*CTCGAG*TAtaaccgcttcatctg |
| hSnd2-$N_S$ | GTGGCCCTC*GAATTC*GAGGAGATCTGCCG | AGGGCCAC*TCTAGA*ttataaccgcttcatc |
| TMEM109-$C_L$ | TAATACGACTCACTATAGG | GAGGGCCAC*GAGCTC*Tctcctcctccacac |
| TMEM109-$C_S$ | TAATACGACTCACTATAGG | GAGGGCCAC*GAGCTC*Tctcctcctccacac |
| Sec61α-$C_S$ | AAGTG*GCTAGC*atggcaatcaaatttc | CCGT*GAATTC*GTgaagagcagggccc |
| Sec61α-$N_L$ | AGCG*CTCGAG*Gatggcaatcaaatttc | GCC*GCTAGC* **TCA**gaagagcagggccc |
| TRAPα-$C_S$ | TAATACGACTCACTATAGG | GGGCCAC*GAGCTC*Cctcatcagatcccac |

Sequences matching cDNA are written in uncapitalized letters, overhangs in capital letters, restriction sites in italic and inserted stop codons in bold.

**Table S20, related to STAR Methods. PCR primers used to create transcription templates.**

| Recombinant cDNA | Vector | Species | Forward primer (5'-3') | Reverse primer (5'-3') | RNA Polymerase |
|---|---|---|---|---|---|
| BCMA-Y13T-OPG2 | pCMV3 | Human | CGCAAATGGGCGGTAGGCGTG | TAGAAGGCACAGTCGAGG | T7 |
| Cytb5OPG2 | pcDNA3 | Human | CGCAAATGGGCGGTAGGCGTG | TAGAAGGCACAGTCGAGG | T7 |
| GypCOPG2 | pGEM4 | Human | GTGGATAACCGTATTACCGCC | CTCTGACGGCAGTTTACGAG | T7 |
| Syt1OPG2 | pcDNA3 | Rat | CGCAAATGGGCGGTAGGCGTG | TAGAAGGCACAGTCGAGG | T7 |
